# Tristetraprolin/ZFP36 regulates the turnover of autoimmune-associated HLA-DQ mRNAs

**DOI:** 10.1101/337907

**Authors:** Laura Pisapia, Russell S. Hamilton, Federica Farina, Vito D’Agostino, Pasquale Barba, Maria Strazzullo, Alessandro Provenzano, Carmen Gianfrani, Giovanna Del Pozzo

## Abstract

We have previously demonstrated that the expression of HLA class II genes is regulated by the binding of a ribonucleoprotein complex that affects the mRNA processing. We identified protein components of a complex binding transcripts encoding the HLA-DR molecule. Here we investigate whether the same RNA binding proteins interact with 3’UTR of mRNAs encoding the HLA-DQ isotype. Specifically, we focused on the HLA-DQ2.5 molecule, expressed on the surface of antigen presenting cells, and representing the main susceptibility factor for celiac disease (CD). This molecule, encoded by HLA-DQA1*05 and HLA-DQB1*02 alleles, presents the antigenic gluten peptides to CD4^+^ T lymphocytes, activating the autoimmune response.

Here, we identified an additional component of the RNP complex, Tristetraprolin (TTP) or ZFP36, a zinc-finger protein, widely described as a factor modulating mRNA stability. TTP shows high affinity binding to 3’UTR of CD-associated HLA-DQA1*05 and HLA-DQB1*02 alleles, in contrast to lower affinity binding to HLA-DQA1*01 and HLA-DQB1*05 non-CD associated alleles. Our *in silico* analysis, confirmed by molecular experiments, demonstrates that TTP specifically modulates the stability of the transcripts associated with celiac disease.

## Introduction

The expression of HLA class II genes is strictly regulated at the transcriptional level by a highly conserved regulatory module, situated 150-300 base pairs upstream of the transcription-initiation site [1]. This regulatory module interacts with several DNA binding factors which in turn bind the HLA class II trans-activator CIITA [2]. We have previously demonstrated that the expression of HLA class II molecules is regulated through a mechanism that coordinates transcription and processing in the context of an “MHCII RNA operon”, a functional unit through which the UTRs of different MHC class II transcripts bind the same RNP complex [3, 4]. We identified two RNA binding proteins, EBP1, the ErbB3 binding protein [5], and NF90, the Nuclear Factor 90 [6], interacting with stem-loop secondary structures at 5’ and 3’ UTRs of HLA-DRA and HLA-DRB1 mRNAs encoding for HLA-DR heterodimer, whose expression is downregulated by the depletion of both proteins. The finding that a specific protein complex regulates the expression of HLA DR transcripts, allows us to extend our investigation to another isotype, HLA-DQ molecule, frequently involved in self-antigen presentation, typical of many autoimmune pathologies.

The HLA-DQ2.5 molecule is strongly associated with celiac disease (CD), a gluten-triggered disorder [7] and type 1 diabetes (T1D) [8], two autoimmune pathologies in which the main genetic predisposing factor is conferred by HLA-DQ2.5 encoding genes. More than 90% of CD patients express the HLA-DQ2.5 molecule, while the majority of remaining affected subjects carry the HLA-DQ8 molecule. In T1D patients, the presence of the HLA-DQ2.5 molecule is less frequent than HLA-DQ8, and 3.5 to 10 % of individuals are affected by co-morbidities of CD and T1D. The surface expression of DQ2 and/or DQ8 molecules, on antigen presenting cells (APC), determines the level of self/gluten-antigen presentation and, as consequence, the magnitude of the pathogenic autoimmune CD4^+^ T cell response leading to organ damage. We recently demonstrated that DQA1*05 and DQB1* 02 genes are more highly expressed than other non-CD-associated alleles in APC from CD patients carrying the DQ2-DR3 genotype in heterozygosis [9]. Furthermore, we found that both DQA1*05 and DQB1*02 mRNAs show a similar stability, although lower with respect to other DQA1*01 and DQB1*05 non-disease associated transcripts. In this study, we aimed to verify if the differential expression of HLA class II transcripts might be ascribed to a specific protein of the ribonucleoprotein complex that preferentially binds to the autoimmune-associated transcripts.

TTP (ZFP36) is a RNA binding protein that preferentially binds to AU-rich regions contained in the 3’UTR of its target genes. TTP functions by destabilizing mRNAs encoding for oncogenes, cytokines (as TNFα) and chemokines involved in the inflammatory processes, by favouring their degradation and/or preventing their efficient translation[10, 11]. Studies in TTP KO mice demonstrate that TTP not only regulates the primary immune cellular response to innate stimuli, but also regulates the expression of transcripts involved in the secondary response of fibroblasts/macrophages to the TNFα stimulation[12, 13]. Here, we show that TTP protein is a component of the MHC operon and is involved in the regulation of the processing of the HLA-DQ genes.

## Results

### TTP binds to CD-associated and not associated transcripts

To analyse the interaction of RNA binding proteins with 3’UTR of DQA1* and DQB1* transcripts we synthetized DNA templates to be used for *in vitro* transcription to obtain the different riboprobes. We used the cDNA of immortalized B cells (B-LCL#5) obtained from a celiac patient carrying DQA1*01-DQA1*05/DQB1*02-DQB1*05 genotype [9] to synthetize the two riboprobes corresponding to 3’UTR of DQA1*05 (3DQA105 riboprobe) and DQB1*02 (3DQB102 riboprobe) mRNAs, encoding the DQ2.5 molecule associated with CD. The cDNA from HOM-2 cell line, carrying the homozygous DQA1*01/ DQB1*05 genotype, was used for the synthesis of two riboprobes corresponding to the 3’UTR of DQA1*01 (3DQA101) and DQB1*05 (3DQB105) mRNAs, encoding DQ5 molecule not associated with disease.

We performed REMSA experiments showing the interaction of 3DQA101 (Figure 1A, lanes 3 and 6) and 3DQA105 (lanes 9 and 12) riboprobes to S100 cytoplasmic extracts of M14 and B-LCL#5, respectively. We also demonstrated that recombinant TTP (rTTP) interact with either 3DQA101 (lanes 4 and 5) and 3DQA105 (lanes 10 and 11) riboprobes. Finally, with an antibody against TTP we carried out a supershift assay. Both 3DQA101 (Figure S1, lane 2) and 3DQA105 (Figure S1 lane 4) riboprobes showed a slower electrophoretic mobility band when anti-TTP was added to the binding reaction in presence of B-LCL#5 extracts.

**Figure 1.**
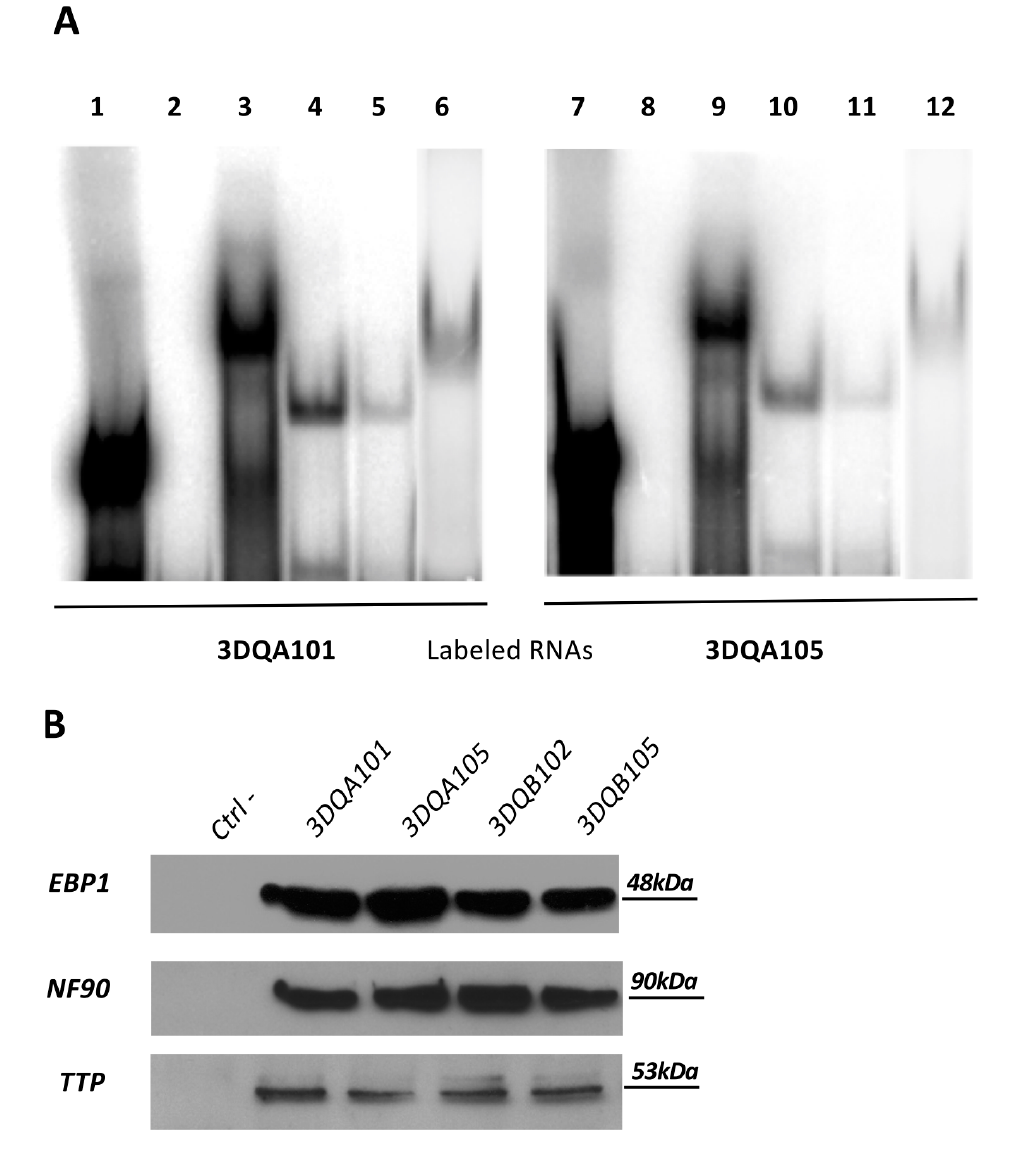
RNA binding proteins interaction. **A.** REMSAs experiments performed using 3DQA101 (lane 1) and 3DQA105 (lane 9) riboprobes. Lanes 2 and 8 show the digestion of riboprobes with T1 RNase. The binding of DQA101 with M14 extract is in lane 3 and with B-LCL#5 extract in lane 6. The binding of 3DQA105 with M14 extract is in lane 9 and with B-LCL#5 extract in lane 12. The interaction of 5 and 1 ug of rTTP with 3DQA101 is showed in lanes 4 and 5 and with 3DQA105 in lanes 10 and 11. **B.** Western blot analysis of biotin pull-down assay carried by using 3DQA101, 3DQA105, 3DQB102 and 3DQB105 riboprobes. The antibodies used for the immunoblot were anti-NF90, anti-TTP, anti-EBP1. Molecular weights are as indicated.

To confirm that this complex includes the same proteins previously identified, we performed a pull-down assay (Figure 1B). *In vitro* transcribed biotinylated riboprobes, 3DQA101, 3DQA105, 3DQB102 and 3DQB105, were incubated with S100 cytoplasmic extract of B-LCL#5 and after pull-down with streptavidin coated beads, proteins interacting with biotinylated riboprobes were detected by western blot analysis. By using anti-NF90, anti-EBP1 and anti-TTP, we demonstrated that all three proteins bind to the 3’UTRs of DQA1* and DQB1* mRNAs. In conclusion, we confirmed that NF90 and EBP1, previously identified as components of the complex interacting with 3’UTRs of DRA and DRB1 mRNA [3, 4], interact with DQ riboprobes while TTP has been for the first time recognized as part of RNP complex binding the HLA-DQA1*01, DQA1*05, DQB1*02 and DQB1*05 mRNAs,

### Analysis of DQA1* and DQB1* 3’UTR sequences

TTP interacts with transcripts via the specific AU-rich AREs in the 3’UTR of target genes and is influenced by the folding of the mRNA itself. We therefore investigated whether differences in the nucleotide sequences may affect RNA folding, the distribution of AU rich elements and their secondary structure accessibility along the 3’UTR. We first compared the nucleotide sequences of the 3’UTRs of DQA1*05 and DQB1*02, CD-associated transcripts, with the 3’UTRs of DQA1*01 and DQB1*05, non-CD-associated transcripts and highlight the differences (Figure 2A,B). In the DQA1* alleles we observed 94.1% identity and a small difference in the overall GC content with the DQA1*01 allele at 0.46 as compared to DQA1*05 at 0.44 (Figure 2A). In the DQB1* alleles, the sequence identity between 3DQB102 and 3DQB105 riboprobes were lower with 92.3%, despite the sequence differences between the alleles the GC content is identical (0.55) (Figure 2B). The sequence differences between alleles are likely to influence the secondary structures of the RNAs, so we performed minimum-free energy (MFE) structure predictions (Figure 2C,D), and mapped on the canonical ARE (AUUUA/AUUUUA) and half ARE (UAUU) motifs associated with TPP binding [11, 14]. A longer length ARE sequence has been defined as WWAUUUAWW [13](W represents A or U), however this motif is not present in the DQA1* or DQB1* 3’UTRs. The canonical ARE and half-ARE motifs are present in both 3DQA101 and 3DQA105 riboprobes, in contrast the ARE motif is only found in the 3DQB102 riboprobe (Figure 2D), suggesting the potential for TTP to bind to CD-associated transcripts. We also performed alignments of the DQA1* and DQB1* alleles with their mouse homologs, H2-Aa and H2-Ab1 (Supplemental Figure 3). The sequence identities of DQA1* and DQB1* with their mouse homologs are <60% and <40% respectively, indicating a low level of conservation of the UTRs across species.

**Figure 2.**
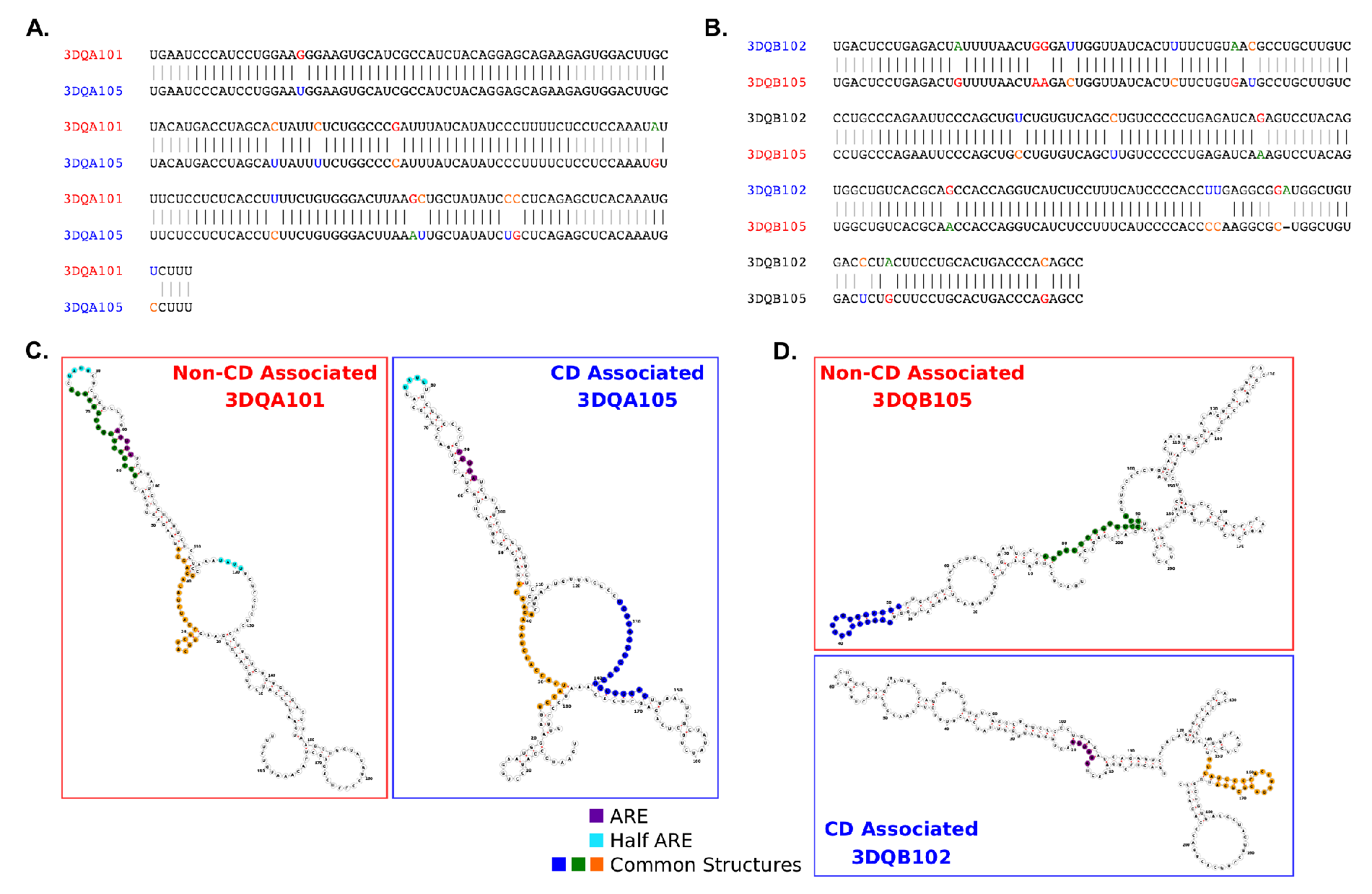
Sequences and structures comparison between 3’UTR of DQA1* and DQB1* genes. **A.** Comparison of the 3’UTR sequences for the DQA1*01 and DQA1*05 alleles indicates a 94.1% sequence identity. There is a slightly higher proportion of GC content in the DQA1*01 allele (0.46) as compared to DQA1*05 (0.44). **B.** The DQB1*05 and DQB1*02 alleles show a lower sequence identity (92.3%), but identical GC proportions (0.55). **C & D.** RNA secondary structure prediction reveals the sequence differences between alleles impacts the minimum free energy folding (RNAfold). Common structural motifs from FOLDALIGN are mapped onto the structures as well as canonical ARE (AUUUA/AUUUUA) and half-ARE motifs (e.g. UAUU).

To assess RNA secondary structure stability between the DQA1* and DQB1* pairs, we calculated ensembles of statistically sampled structures using Sfold [15] (Supplementary Figure 2). The ensembles represent the full range of structures a sequence can adopt, not just those with the lowest MFE, highlighting substructures that remain constant across all structure conformations. MFE prediction makes the assumptions that there is a single rigid structure and that the energy parameters are complete. However, it is likely that messenger RNAs are more dynamic and can adopt multiple conformations depending on their environment and presence of RNA binding proteins. Across the predicted ensemble structures the conserved substructures co-localise with the ARE and half-ARE sequence motif positions in the 3DQA101, 3DQA105 and 3DQB105 mRNAs. The 3DQB102 ARE is outside of a predicted stable substructure, indicating it is likely to be more flexible and adopt a range of structural conformations. The proportion of the 3DQB102 structure with predicted conserved substructures between ensembles and minimum-free energy predictions, and those from FOLDALIGN (see below), is lower than in the other alleles and importantly, does not contain AU-rich elements indicating that TTP may bind in regions with low structural stability.

We also performed structural motif searches between pairs of 3’UTRs with FOLDALIGN [16] to determine if there are any common structural motifs (defined by sequence and secondary structure) that could confer an RNA Operon-like module as previously found [4]. The presence of a common motif present in different structures could indicate a common RNA-binding protein site. The identified common structural motifs were then mapped onto the structures (Figure 2C,D). We found structures common to the DQA1* and DQB1* pairs and we hypothesis this could enable them to co-bind in an “RNA Operon” as we previously demonstrated for HLA-DR alleles [4].

We performed a more extensive AU rich sequence motif search to determine if there are further possible, weaker TPP binding sites in riboprobes and whether there are differences between the CD and non-CD associated allele pairs. Our kmer analysis at a range of motif lengths (di, tri, tetra, penta and hexa nucleotides) shows a clear pattern of enrichment of AU containing motifs in the CD associated alleles as compared to the non-disease associated across all kmer lengths (Figure 3A). Mapping the AU rich kmers onto the riboprobe sequences revealed the CD associated alleles contain more AU rich sites located across the full length of the sequence and therefore are likely to confer more TPP binding through these AU motifs. Individual AU rich kmers are also more commonly seen in clusters together in close proximity in the CD associated alleles than those non-disease associated (Figure 3B).

The secondary structure context of the AU rich kmers will influence the ability of TPP to bind in a sequence specific manner. We therefore utilised RNAplfold to calculate a secondary structure accessibility score, indicating the probability of whether the RNA is likely to be single stranded along its length. In the DQA1* alleles the AU motifs are more prevalent in the CD associated allele, however they tend to be localised in regions of a lower probability of being single stranded than the non-disease associated allele. In DQB1* the AU motifs in the CD-associated alleles are not only more common but also colocalised with single stranded regions, in contrast the AU motifs in the non-disease allele are not in single stranded regions.

In conclusion, our analysis predicts that TTP binds to HLA-DQA1 and HLA-DQB1 mRNAs and that the secondary structures adopted by the alleles facilitate the interaction of TTP with the 3’UTR of CD-associated mRNAs. According to the canonical role of TTP, this binding should confer lower stability to transcripts respect to non-CD associated mRNAs.

**Figure 3.**
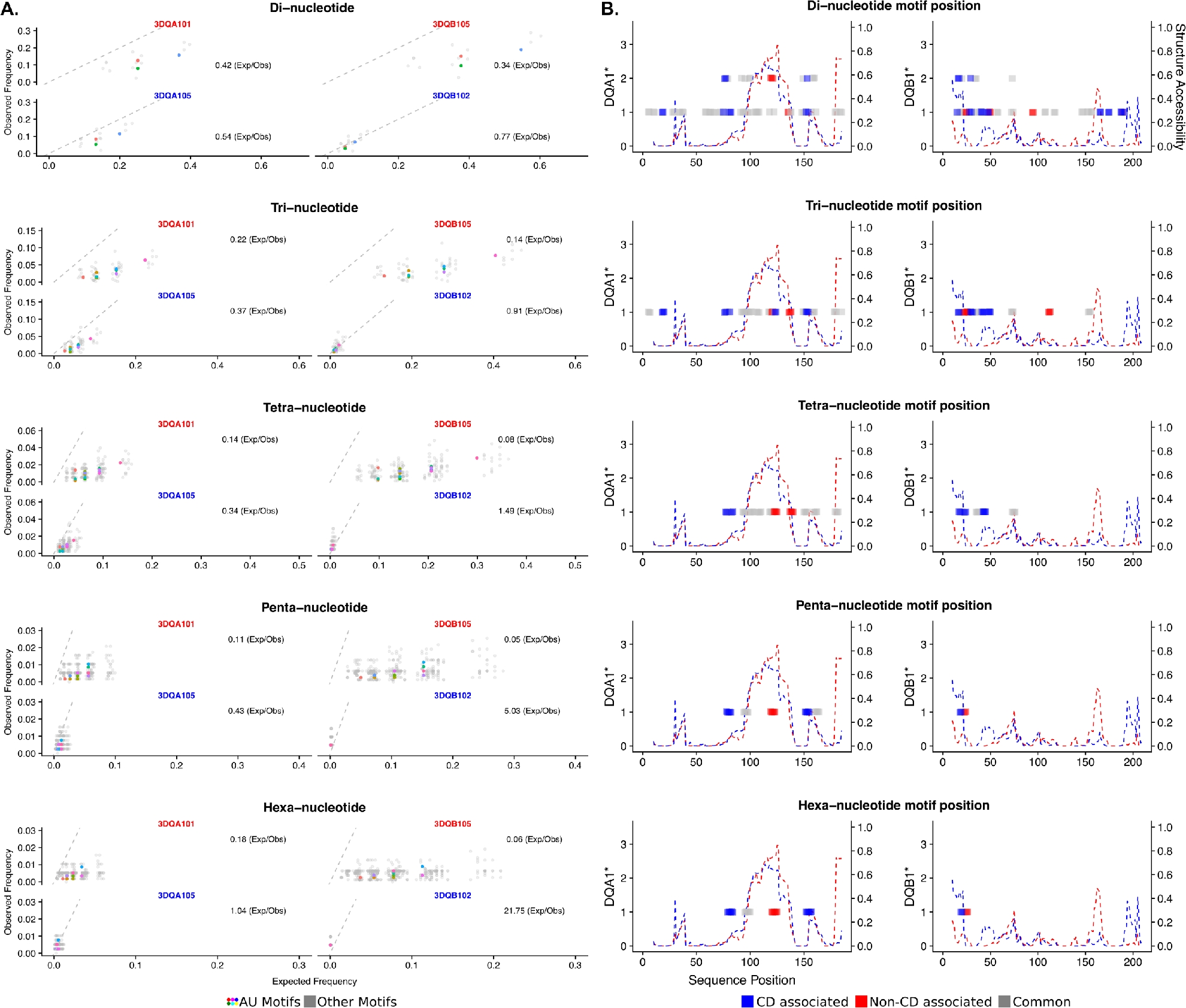
AU-rich motif analysis of the DQA1* and DBQ1* genes reveals key differences. **A.** All possible di-, tri-, tetra-, penta- and hexa-mers are assessed for their observed frequencies against their expected frequencies. AU-rich motifs are colored, non-AU rich in grey. All AU-rich kmer motifs have higher enrichments in the CD associated DQA1*05 and DQB1*02 alleles as compared to the non-CD associated DQA1*01 and DQB1*05 alleles. **B.** Mapping the AU-rich motifs to their position within the 3’UTR sequences reveals a clear enrichment in AU-motifs throughout the CD associated alleles DQA1*05 and DQB1*02 (Blue) as compared to the non-CD associated DQA1*01 and DQB1*05 alleles (Red). Nested motifs are indicated on the y-axis, for example two overlapping motifs will show a score of 2. A structure accessibility score indicates the likelihood the RNA single stranded, an absence of base-pairing, across its length. A score of 1.0 indicates the structure is single stranded.

### Knockdown and overexpression of TTP affects the expression level of DQA1 and DQB1 transcripts

We previously showed a differential expression of CD-associated transcripts, DQA1*05 and DQB1*02, and their lower half-lives in respect to non-CD associated DQ transcripts in B-LCL#5 cells [9]. In order to investigate if the TTP protein is responsible for the DQ mRNAs turnover, we depleted the protein by specific siRNAs transfection in M14 and B-LCL#5 cell lines. The knockdown of TTP was first assayed in M14, 48 h after transfection by western blot with anti-TTP (Figure 4A) in which we observed 80% depletion. When we measured the mRNA amount by qRT-PCR, we found 2 to 3 fold increase of DRA, DRB1* and DQA1* transcripts (Figure 4B). As a control of the knockdown specificity, we showed no variations in the expression of two groups of genes related to our system, the HLA-A, B and C class I genes and CIITA, the HLA class II transcriptional activator. In parallel, we performed TTP knock-down in B-LCL#5 by specific siRNA nucleofection and confirmed the protein depletion after 48 h by western blot (Figure 5A).

**Figure 4.**
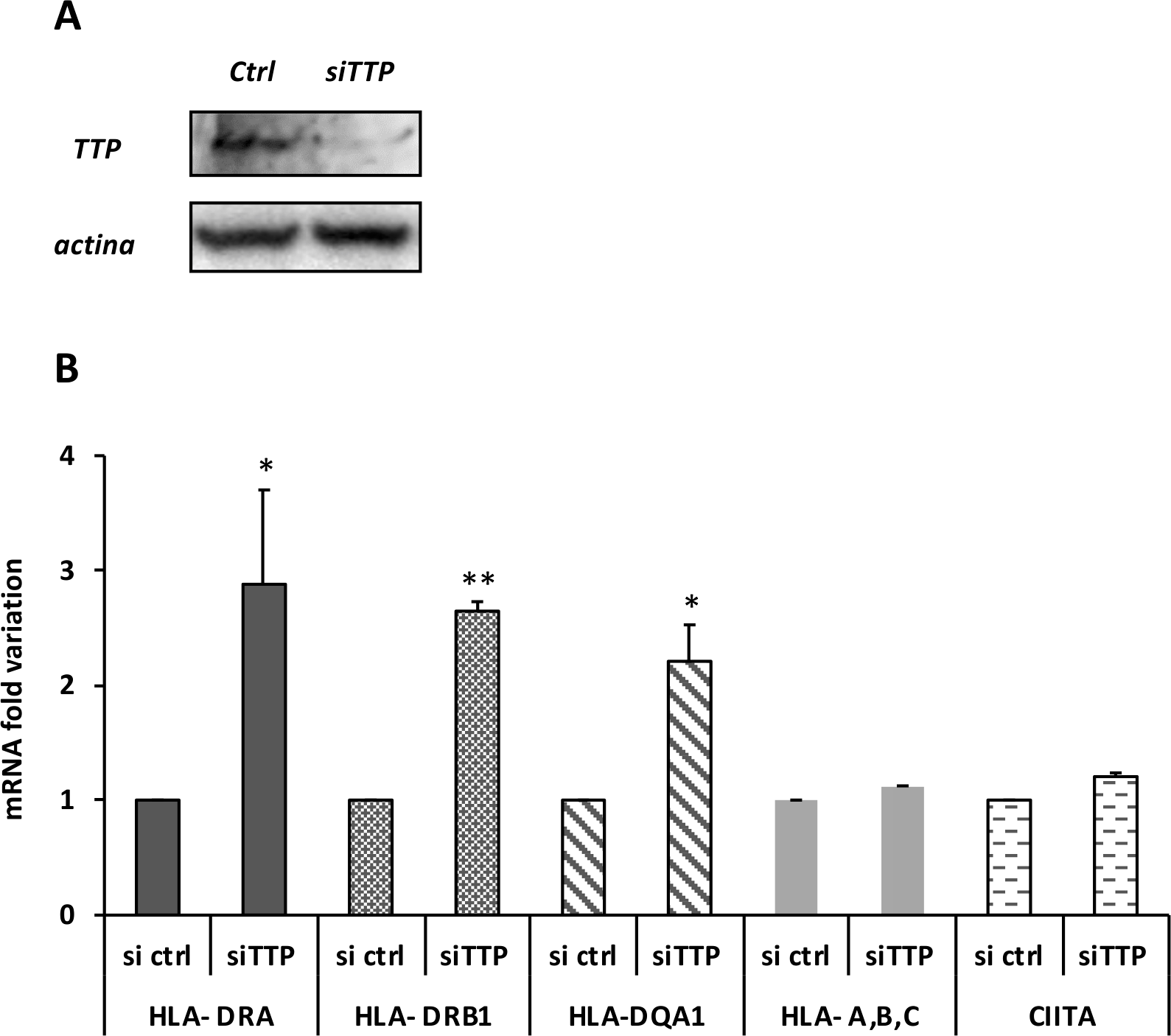
TTP knockdown in M14. **A.** western blot performed with anti-TTP for the assessment of protein depletion 48 h after silencing with siCtlr or siTTP **B.** Fold variation of DRA, DRB1, DQA1 mRNAs. HLA-a,b,c are the class I mRNAs, CIITA is the HLA class II transcriptional transactivator.

The TTP knockdown affects the phenotype of our cells and, by flow cytometry analysis, we observed higher HLA-DQ surface expression (Figure 5B). To evaluate whether the DQ increase corresponds to the HLA-DQ mRNAs variation we carried out qRT-PCR. We found a 2.7 and 2.9-fold increase of DRA and DRB1* mRNAs, respectively, while total DQA1* and DQB1* mRNAs show 3-fold increment compared to the control (siCtlr) and respect to the HLA class I mRNAs (Figure 5C). To investigate if TTP affects the expression of each mRNAs, we specifically quantified DQA1*01, DQA1*05, DQB1*02 and DQB1*05 mRNAs. The experiment confirmed that DQA1*05 and DQB1*02 have a higher expression than DQA1*01 and DQB1*05 (Figure 5C) as previously demonstrated [9]. However, when we compared the amount of different messengers after TTP knockdown, we observed that DQA1*05 and DQB1*02 mRNAs showed a greater and significant increment (3,5-fold increase), respect to DQA1*01 and DQB1*05 mRNA (2-fold increase, Figure 5D). In conclusion, we demonstrate that the depletion of TTP protein, interacting with 3’UTR, determines an increment of the CD-associated DQA1*05 and DQB1*02 mRNAs greater than that of the non-CD-associated DQA1*01 and DQB1*05 mRNAs.

**Figure 5.**
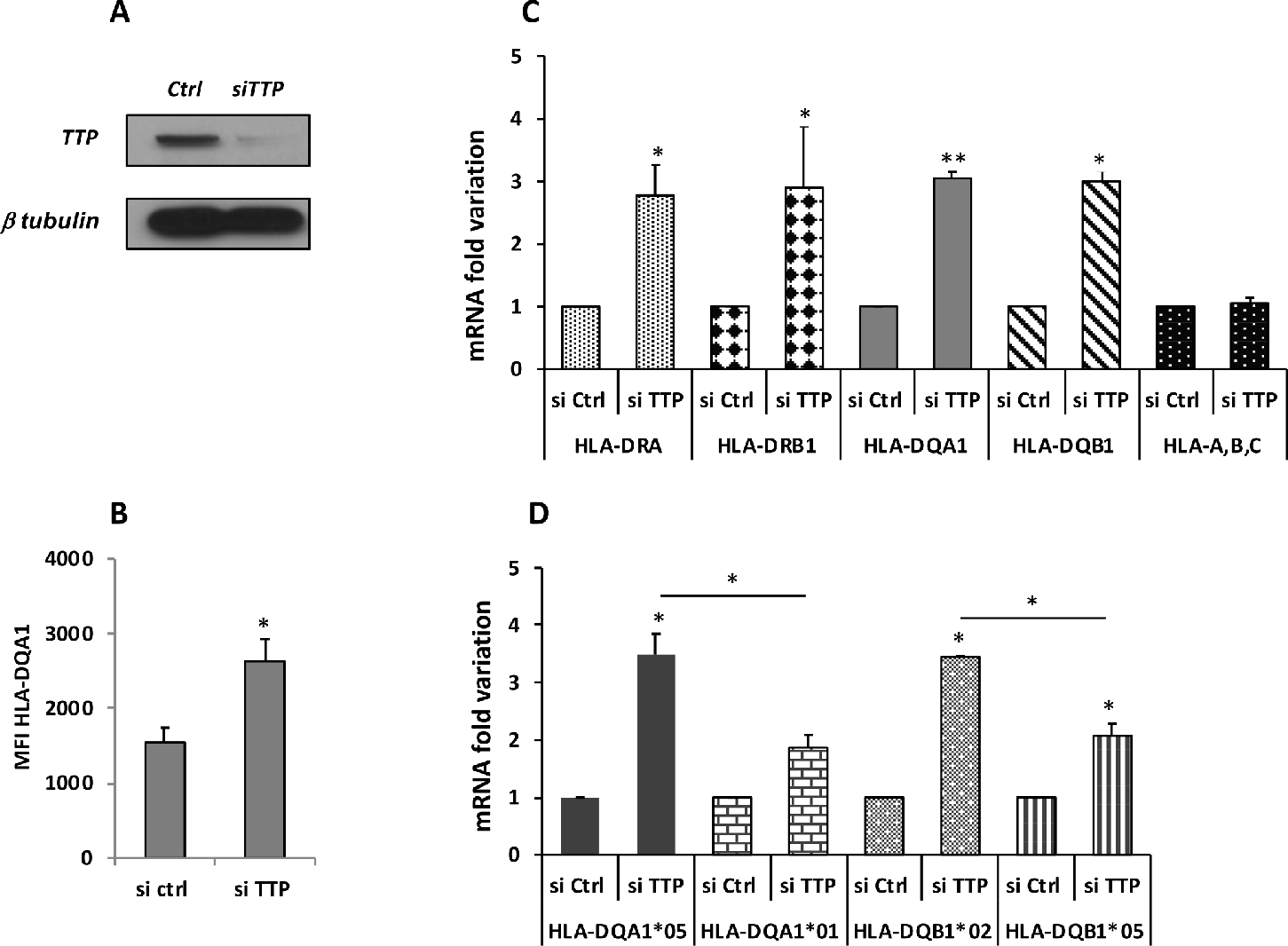
TTP knockdown in B-LCL#5. **A.** Western blot performed with anti-TTP for the assessment of protein depletion 48 h after silencing with siCtlr or siTTP nucleofection. **B.** Cytofluorimetric analysis of HLA-DQ surface expression, reported as fold change of MFI (Mean Fluorescence Intensity). **C.** Fold variation of DRA, DRB1, DQA1, DQB1 and HLA-A,B,C (class I) mRNAs. **D.** Fold variation of DQA1*01, DQA1*05, DQB1*02 and DQB1*05 mRNAs.

We also performed the overexpression of TTP protein by nucleofection of cDNA in BLCL#5 and assessed the protein overexpression after 48 h by western blot (Figure 6A). As a positive control of TTP overexpression, we evaluated, by qRT-PCR, the amount of TNFα transcript, and showed a 17 fold decrease (Figure 6B), as expected. Then we quantified the HLA-DQ alleles levels and demonstrated 17 fold decrease of DQA1*05 and 4,5 fold decrease of DQB1*02, as expected, while surprisingly, TTP overexpression causes the increase of DQA1*01 and DQB1*05 (4,5 and 6,7 fold variation, respectively) mRNAs. In conclusion, the ectopic expression of TTP protein differently influences the amount of transcripts, producing a reduction of DQA1*05 and DQB1*02 and an increment of DQA1*01 and DQB1*05 mRNAs.

**Figure 6.**
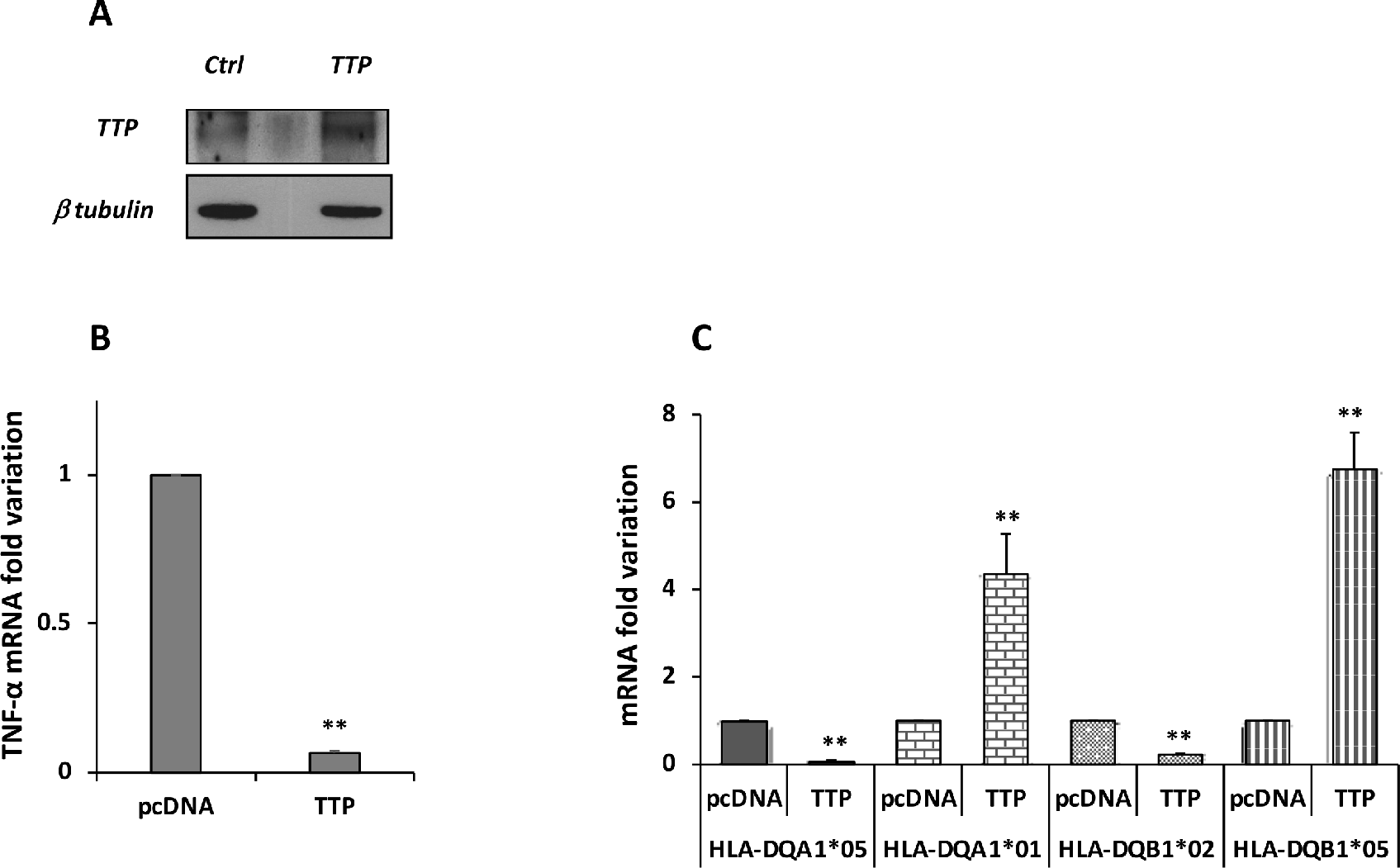
TTP overexpression in B-LCL#5. **A.** Western blot performed with anti-TTP for the assessment of protein ectopic expression 48 h after nucleofection. **B.** Fold variation of TNFα mRNA. **C.** Fold variation of DQA1*01, DQA1*05, DQB1*02 and DQB1*05 mRNAs.

## Discussion

HLA class II molecules are expressed on the surface of professional antigen presenting cells (APC) such as monocytes, dendritic cells, B lymphocytes, macrophages, and of non-professional APC, such as cancer cells, with the role of presenting antigen epitopes to CD4^+^ T cells. The magnitude of the CD4^+^ T cell activation and proliferation is related not only to the nature of cognate epitopes, but also to the amount of the antigen-HLA class II complexes expressed on the surface of APC [17]. In this respect, the expression level of HLA class II molecules that restricts the antigenic responses is very important, particularly in autoimmune diseases as the HLA class II encoding genes represent the main genetic risk factor associated with these pathologies.

Previously, we demonstrated similar amounts of DQ2.5 molecules in APC of CD patients either heterozygous or homozygous for DQ2.5 risk genes [9]. As consequence, DQ2.5 heterozygous or homozygous APC induced a similar strength of functional immune response by gluten-reactive CD4^+^ T cells. The equivalent surface density of DQ2.5 heterodimer on cells from homozygous and heterozygous CD patients was due to the marked expression of DQA1*05 and DQB1*02 risk genes, in respect to the reduced expression of non-disease associated DQA1*01 and DQB1*05 alleles, in DQ2.5 heterozygous genotype. This difference in the quantity of transcripts might be explained by haplotype-specific transcriptional regulation or allele-specific mRNA processing [17]. Indeed, the transcripts of both CD-associated DQA1*05 and DQB1*02 genes showed a similar decay kinetic (3hr half-live), but much lower in respect to non-CD-associated DQA1*01, DQB1*03 and DQB1*05 mRNAs (4h half-live) [9]. This difference in the transcript turnover led us to hypothesize that a protein complex might differentially bind the autoimmune-associated messengers, and therefore modulate their decay.

The analysis of 3’UTR of CD-associated DQA1*05 and DQB1*02, and non-CD associated DQA1*01 and DQB1*05 alleles revealed several differences in GC content and in canonical ARE (AUUUA) and half ARE (UAUU) motifs in addition to base differences that influence the secondary structures and binding sites of the RNAs. The predicted RNA structures show no differences in the presences motifs containing substructures between DQA1* mRNAs, but lower structural stability for DQB1*02. The lower stability of this transcript influences the halflife of DQA1*05, as the turnover of two mRNA is co-regulated. In other words, the amount of a messenger establishes the quantity of its partner messenger coupled in the same RNP complex [4, 17]. REMSA confirmed the *in silico* findings, revealing the interaction of cytoplasmic extracts with 3’UTR of DQA1*01 and DQA1*05 transcripts. Importantly, this experiment indicates that the RNA binding proteins included in the complex are constitutively expressed by either professional (B-LCL) and/or non-professional antigen presenting cells (M14). Moreover, in these experimental conditions, the 3’UTR of DQB1*02 and DQB1*05 RNAs do not show binding with cytoplasmic extracts. The absence of binding with beta riboprobes may be caused by the radiolabelling of the 3’UTRs affecting their secondary structures. For this reason, we investigated the RNA-protein interaction by an end-labelled desthiobiotin based pull-down method that, does not interfere with the RNA structure. Indeed, following the efficient enrichment of the protein-RNA complexes, we assessed by immunoblot, the interaction of two RNA binding proteins, EBP1 and NF90, previously identified in the complex with of DRA and DRB1* 3’UTR [3, 4], with four 3DQA105, 3DQA101, 3DQB102 and 3DQB105 riboprobes analysed in this work.

To investigate the different turnover rates of the mRNAs, we evaluated the interaction of TTP (ZNFP36), a zinc finger protein interacting with AU-rich elements (ARE)-containing mRNAs promoting rapid cytoplasmic decay through acceleration of the deadenylation rate [18]. We assessed the interaction of recombinant TTP to the 3’UTR of all transcripts either by REMSA and pull-down experiment. In addition, to verify if TTP protein affects the stability, we performed its depletion by specific siRNA, and observed, as expected, an increase of HLA class II mRNAs, in both cell lines, while HLA class I and CIITA mRNAs were unaffected. Notably, in B-LCL#5, we found that TTP depletion results in a 3.5-fold increase of DQA1*05 and DQB1*02 mRNAs and 2-fold increase of DQA1*01 and DQB1*05 mRNAs. This result clearly indicates that the binding of TTP destabilizes CD-associated messengers more than non-CD associated ones. Moreover when we performed TTP overexpression, we found a decreased level of DQA1*05 and DQB1*02 mRNA similar to TNFα and surprisingly an increase of DQA1*01 and DQB1*05 mRNAs. This results might be explained with the need of APC to maintains stable the concentration of each HLA class II isotype, in this case DQ molecules, on cell surface. An interplay between transcriptional and post-transcriptional would explain the balanced expression of alleles of the same isotype.

Tristetraprolin has previously been shown to bind via canonical ARE (AUUUA) motifs, present in the upstream region of 3DQA101 and 3DQA105 riboprobes, in addition to the ARE motifs there are also half-ARE (UAUU) motifs present in the 3DQA101 (two copies) and 3DQA105 riboprobes (Figure 1). More recently, TTP has been demonstrated to bind to AU-rich sequences [14] [11], and through our sequence analysis we have demonstrated there is an enrichment of AU-rich motifs with TPP binding potential in the CD-associated alleles as compared to the non-CD associated. These motifs are located throughout the sequences of the CD-associated alleles in contrast the non-CD-associated alleles where they form a cluster between positions 90-160 (approximate) (Figure 2). The sequence and AU-rich motif distribution differences between the DQA1 and DQB1 alleles is also likely to have an impact on the secondary structures of the RNAs and their accessibility to TPP binding. Our analysis of the ensembles of predicted structures indicates that, in addition to the sequence motifs, there are differences in the structure stabilities contributing to the observed differences of TPP binding affinities (Supplemental Figure 2). Across all the secondary structure conformations (ensembles) that can be adopted by each of the RNA alleles, we observed that ARE motifs are co-located with the substructures present across all ensemble structures, thus indicating that they are present in the more stable sections of the structures. In the case of 3DQB102, there is an ARE not found in the 3DQB105 riboprobe, indicating that TPP binding may higher affinity in 3DQB102, although this ARE is outside of the more stable substructures. In conclusion, our bioinformatics analysis confirms that CD-associated mRNAs can strongly interact with TTP through their 3’UTR, conferring a more rapid mRNA turnover.

We also investigated mRNAs controlling inflammation in mouse, where is reported that no significant binding of TTP has been detected to H2-Aa and H2Ab1 mRNA, the mouse HLA-DQ-homologous transcripts [13]. However there is low sequence identity between human and mouse HLA-DQ 3’UTRs (See Figure S3). Our results therefore provide evidence that the function of TTP as an RNA-binding protein is more context-dependent than previously described in literature and the canonical binding site is present in stable as well as unstable TTP-bound mRNAs.

In addition, it has been demonstrated in LPS activated macrophages that there is a strong impact of TTP-dependent mRNA degradation on the transcriptome, during the early resolution phase of inflammation than during the onset of inflammation [13]. Inflammation is linked to the adaptive immune response since, following HLA-antigen complexes recognition, proliferating CD4^+^ T cells activates macrophages through cell-to-cell contact and IFN-γ secretion. We propose that TTP-mediated regulation of the HLA-DQ2 mRNAs stability occurs during the resolution phase of inflammatory response. TTP interaction with these transcripts may counterbalance the high transcriptional expression of DQ2 genes and modulate the CD- and T1D-related antigen presentation.

## Materials and Methods

### Cell lines

M14 human melanoma and human HOM cell lines (DQA1*01/DQB1*05 genotype) were obtained by ECACC. B-LCL#5 was EBV-transformed B lymphoblastoid cell lines immortalized from PBMCs of celiac patients carrying DQA1*01-DQA1*05/DQB1*02-DQB1*05 genotype [9]. All cell lines were cultured in RPMI 1640 medium supplemented with 10% Foetal Calf Serum (FCS).

### DNA templates synthesis and mRNA quantification

The sequences of specific HLA alleles were obtained from the HLA database (http://www.ebi.ac.uk/ipd/imgt/hla/index.html) and the primers used for PCR and qRT-PCR were synthesized by Eurofins and listed in Table 1. The template for 3DQA105 riboprobe synthesis was previously prepared [3], while the others obtained from retro-transcribed RNA. Specifically, we used cDNA from B-LCL#1 to obtain 3DB102 template and cDNA from HOM cell to synthetize 3DQA101 and DQB105 templates. Total RNA was prepared with the AurumTM Total RNA kit (BIORAD), and 1 ug of RNA was used for reverse transcriptase reactions, performed using an iScriptTM cDNA Synthesis kit (BIORAD). To quantify specific transcripts, we performed qRT-PCR using the Quanti Tect SYBR Green PCR Kit (BIORAD) through the DNA Engine Opticon Real-Time PCR Detection System (BIORAD). Each reaction was run in triplicate in the presence of 0.2 mM primers. The relative amount of specific transcripts was calculated by the comparative cycle threshold method [19], and GAPDH and β-actin transcripts were used for normalization. All results shown are the mean of at least three independent experiments. Statistical analysis was performed using the unpaired Student’s t-test with two-tailed distribution and assuming two samples equal variance parameters. In the figures, a single asterisk corresponds to *p*< 0.05 and double asterisks correspond to *p*< 0.01.

**Table 1.**
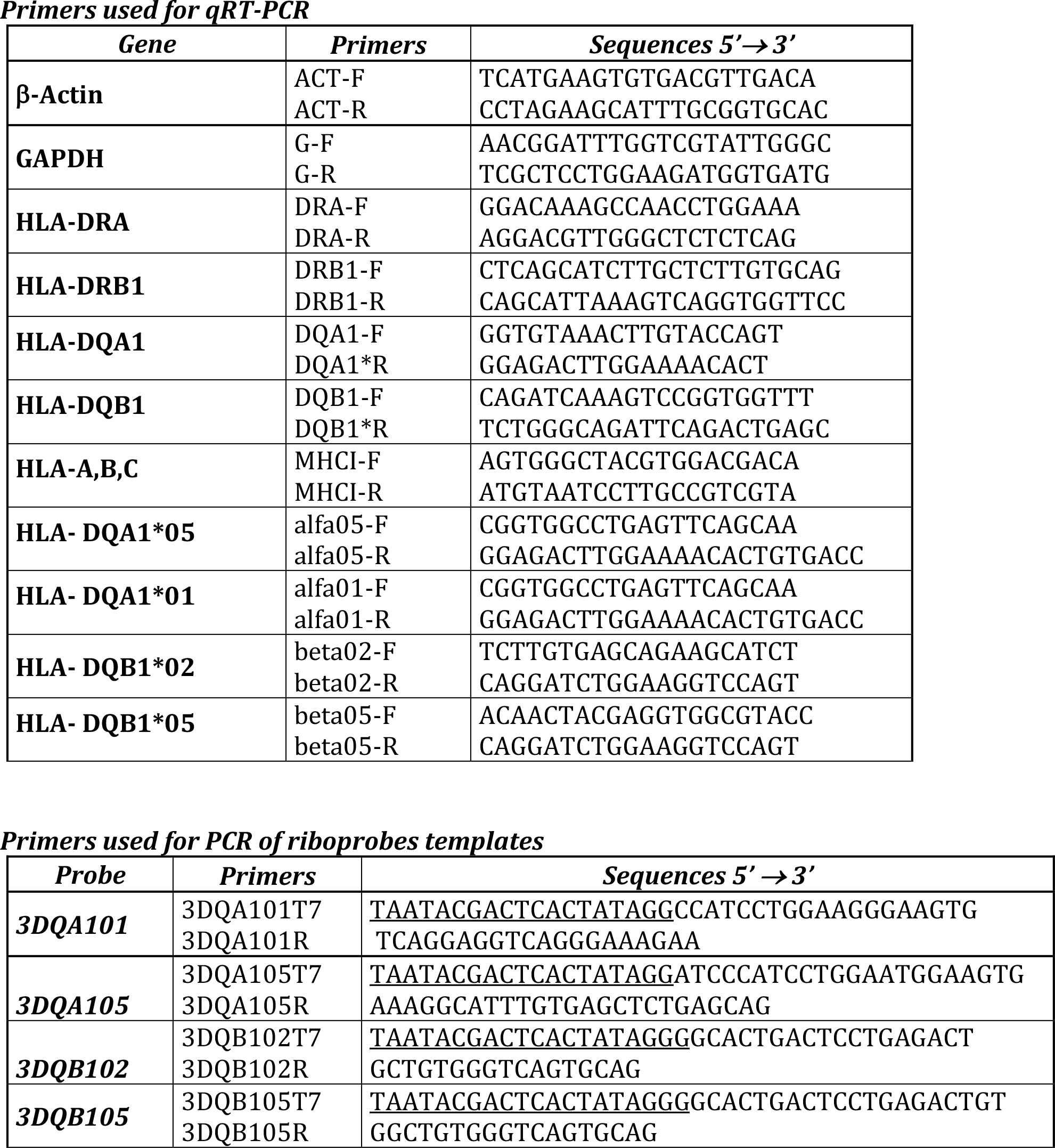
Primers used for qRT-PCR.

### RNA electrophoretic mobility shift assay (REMSA) and pull-down

The riboprobes synthesis and REMSA were performed according to the published protocol [3, 4]. Briefly, the transcription reactions were performed using T7 *in vitro* transcription system (Ambion) in presence of [^32^P] UTP and riboprobes obtained were used in binding experiments with M14 and B-LCL#5 S100 extract. TTP recombinant proteins was produced as described [20]. The antibody supershift assay was carried out with 4 μg of specific polyclonal rabbit anti-TTP that were pre-incubated with cell extracts prior to the addition of riboprobes.

For pull-down experiments, riboprobes were end-labeled with desthiobiotin cytidine and used in binding experiments with 60 ug of B-LCL#5 cytoplasmic extract with the Thermo Scientific Pierce Magnetic RNA-protein pull down kit. The riboprobe used as negative controls was the 3’UTR of androgen receptor RNA poly(A) _25_ RNA, provided by the kit.

Desthiobiotinylated target RNAs bound to proteins were captured using streptavidin magnetic beads and following washing and elution, the proteins interacting with RNA were separated by SDS-PAGE and analysed by western blot. We used three different antibodies: anti-DRBP76 (anti-double stranded RNA binding protein 76 or anti-NF90) antibody (BD Biosciences), N-terminus anti-EBP1 (Abcam), anti-TTP (Tristetraprolin, Santa Cruz Biotechnology), to reveal the presence of proteins in the RNP complex binding 3’UTR.

### TTP protein expression, gene silencing and phenotype analysis

The plasmid for recombinant wild-type (AA) His-tagged TTP proteins (kindly provided by Dr. Tiedje) have been used to transfect HEK293T cells and protein purification using nickel-chelate agarose beads and following a protocol already described [20]. After imidazole elution, samples were dialyzed and stored at −80°C in a solution made of 20 mM HEPES pH 8, 100 mM NaCl, 3 mM MgCl_2_, 8% Glycerol.

For TTP depletion we performed gene silencing using a pool of 4 different siRNA provided by Santa Cruz Biothecnology. We used HiPerFect Transfection Reagent (QIAGEN) to transfect M14 cells and 5×10^5^ cells were harvested after 48 h for RNA extraction and after 24 h for protein extraction. Either TTP depletion and overexpression in B-LCL#5 were performed by using nucleofector technology with Lonza kit. 5×10^5^ cells were transfected with siRNA pool or with pcDNA carrying TTP cDNA used for the synthesis of rTTP. The cells were harvested after 48 hrs either for protein extraction, flow cytometry analysis and RNA preparation. TTP depletion and overexpression were assessed by western blot using an anti-TTP antibody. The HLA-DQ cell surface expression was performed by cytofluorimetric analysis using the FACSAria III and DIVA software with FITC mouse anti-human HLA-DQ antibody (BD Biosciences). The quantitation of specific transcripts was performed by qRT-PCR as described in [9], using primers listed in Table 1.

### Bioinformatics Analysis

Sequence alignments and identities were calculated using the global Needleman-Wunch algorithm [21]. Minimum Free Energy (MFE) RNA secondary structures were predicted using RNAfold [22] from single sequences. Structure accessibility probability scores were calculated using RNAplfold,[22] using a window size of 75 and mean structure score for fragments of 10 nt (RNAplfold -T 37 -W 75 -u 10). [13]These parameters are similar to those used in the TPP Atlas describing transcriptome-wide TPP binding. In addition to MFE structure calculations, an ensemble of statistically sampled structures was generated using Sfold [15], for each riboprobe sequence indicating the most structured parts of the molecules common to the predicted ensemble clusters. To identify common structural motifs between the DQA1 and DQB1 sequences we ran Foldalign [23], in a pairwise all-against-all search. Bracket notation secondary structure from RNAfold were rendered into images with selected motifs mapped onto them using FORNA [24].

AU rich motif mapping and enrichment analysis was performed with custom scripts, freely available at https://github.com/darogan/2018PisapiaDelPozzo including R code to recreate Figure 3. All possible kmers, for lengths 2, 3, 4, 5 and 6, were created for each of the riboprobe sequences, and the observed versus expected ratios calculated. Subsets of AU rich motifs are highlighted in the plots. We then mapped the same AU motifs onto each of the riboprobe sequences. Nested structures, i.e. overlapping, are indicated by the score on the y-axis of Figure 3.

## Acknowledgements

This work was supported by CNR-DSB Progetto Bandiera “InterOmics” 2017 to CG, MS and GDP. RSH is funded by the Centre for Trophoblast Research, University of Cambridge.

## Supplementary Information

**Figure S1.**
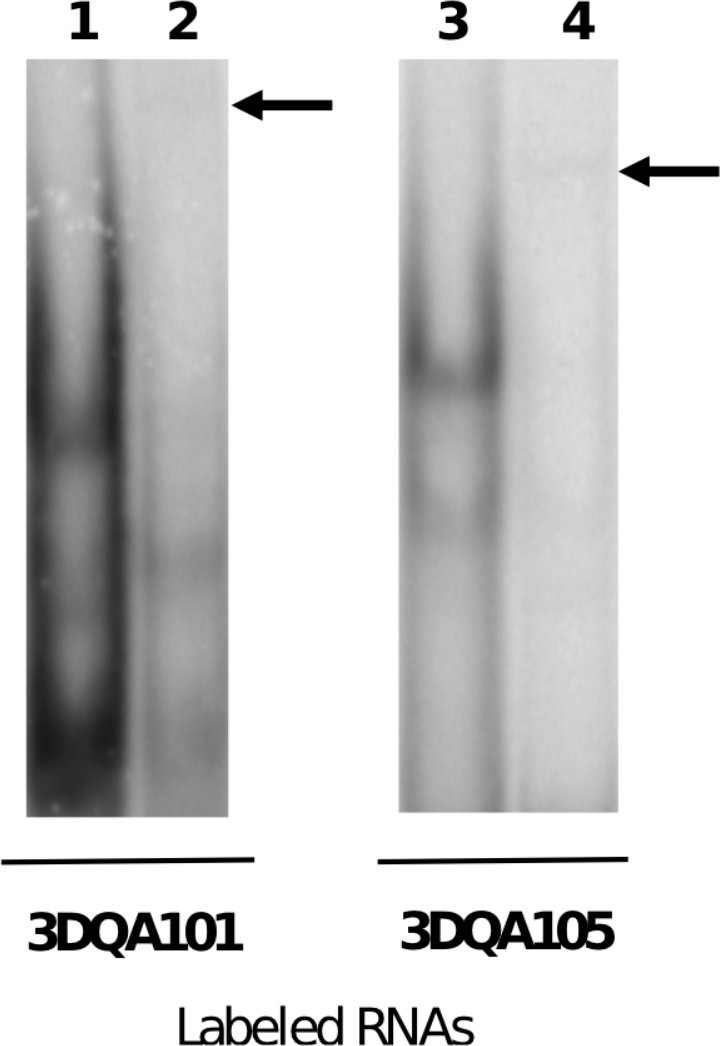
REMSA showing RNA supershift. The interaction between B-LCL#5 extracts with 3DQA101 (lane 1) and 3DQA105 (lane 9) riboprobes were performed in presence of the anti-TTP antibody. The arrows show the retarded band in presence of antibody (lane 2 and 4, respectively)

**Figure S2.**
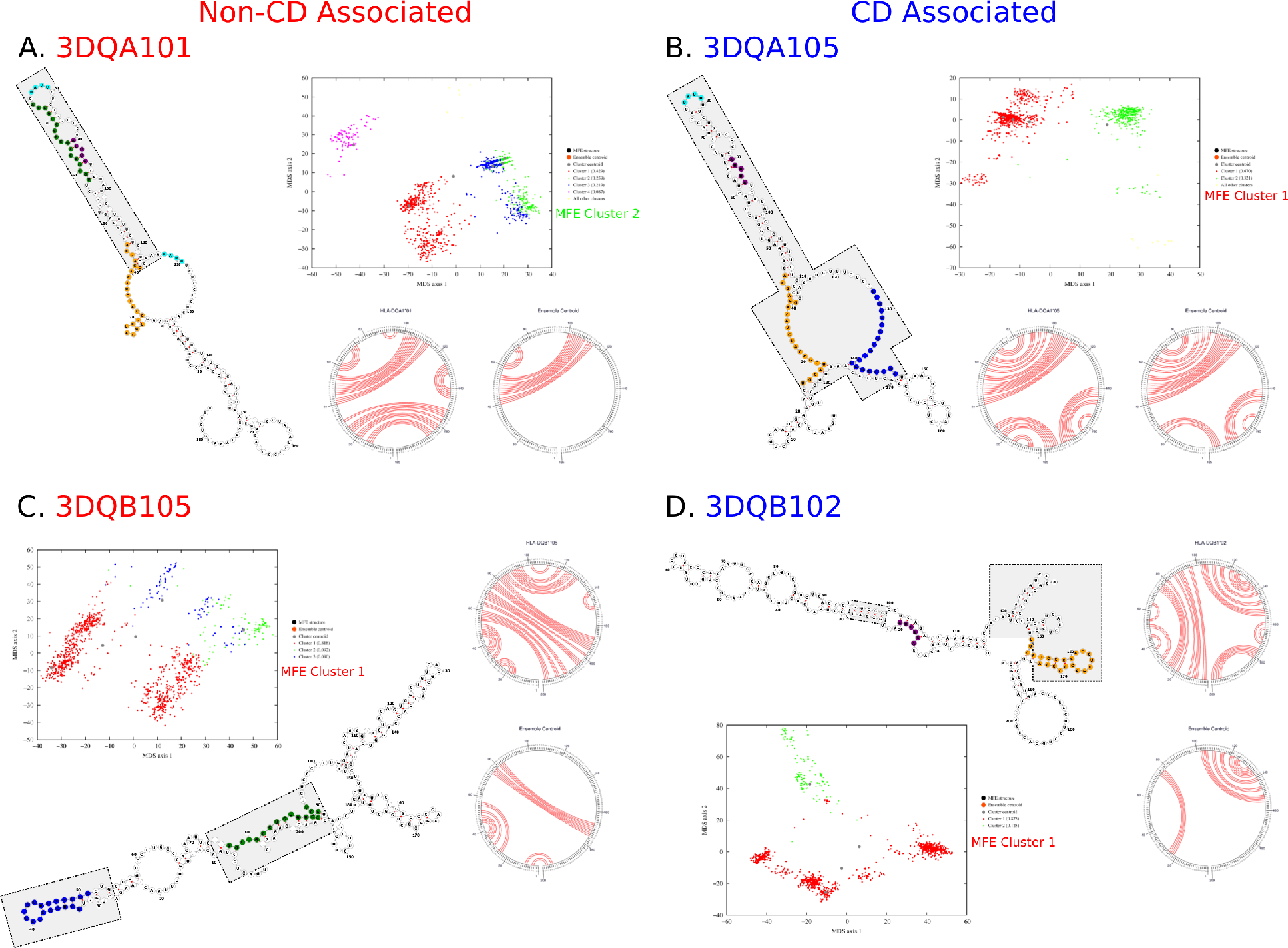
Sfold Structure Comparison of 3’UTR of DQA1* and DQB1*. Each of the riboprobe sequences was analysed with Sfold to predict statistical ensembles of structures. The ensembles permit many different conformations of the structures and clustering indicating the most likely conformations adopted by each sequence. The minimum-free energy (MFE) structure is also predicted and its parent cluster identified. **A.** 3DQA101, **B.** 3DQA105, **C.** 3DQB105 and **D.** 3DQB102 show the ensemble structures to contain fewer conserved base pairings (grey boxes) than the MFE structure indicating the structure is more dynamic/flexible outside of these regions. This is particularly evident in **D.** 3DQB102 where there is little conservation of base pairings in the large stem as seen in the other structures.

**Figure S3.**
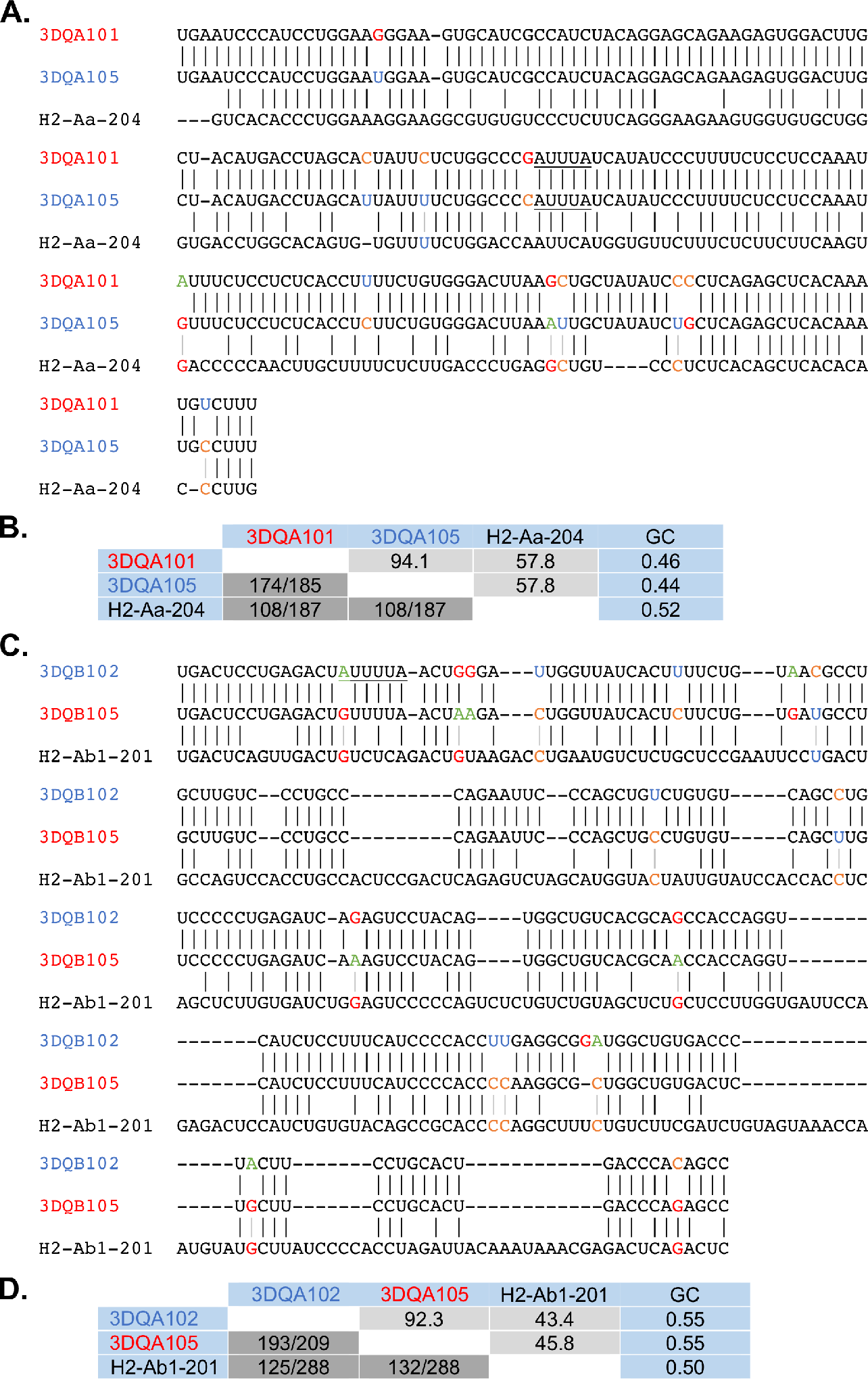
Sequence Alignment for human and mouse homologs for DQA1* and DQB1* alleles. The sequence alignments for the human DQA1* and DQB1* 3’UTR sequences in Figure 1A and B have been extended to include the mouse homologs H2-Aa and H2-Ab1. **A.** The DQA1* alignment for the human alleles and mouse homolog show high sequence identity between the human alleles at 94.1% but a much lower identity between the human and mouse sequences (<60%) as summarised in **B. C.** The DQB1* alignment for the human alleles and mouse homolog show a low sequence identity of <45% between species but >90% for the human sequences as summarised in **D.** The low level of conservation between human and mouse (<60% for DQA and <45%DQB) will result in different locations of AU rich motifs and affect their presentation in single stranded RNA structure regions.

