## Supplementary figures and images for "Tristetraprolin/ZFP36 regulates the turnover of autoimmune-associated HLA-DQ mRNAs"

### Supplemental Fig 2

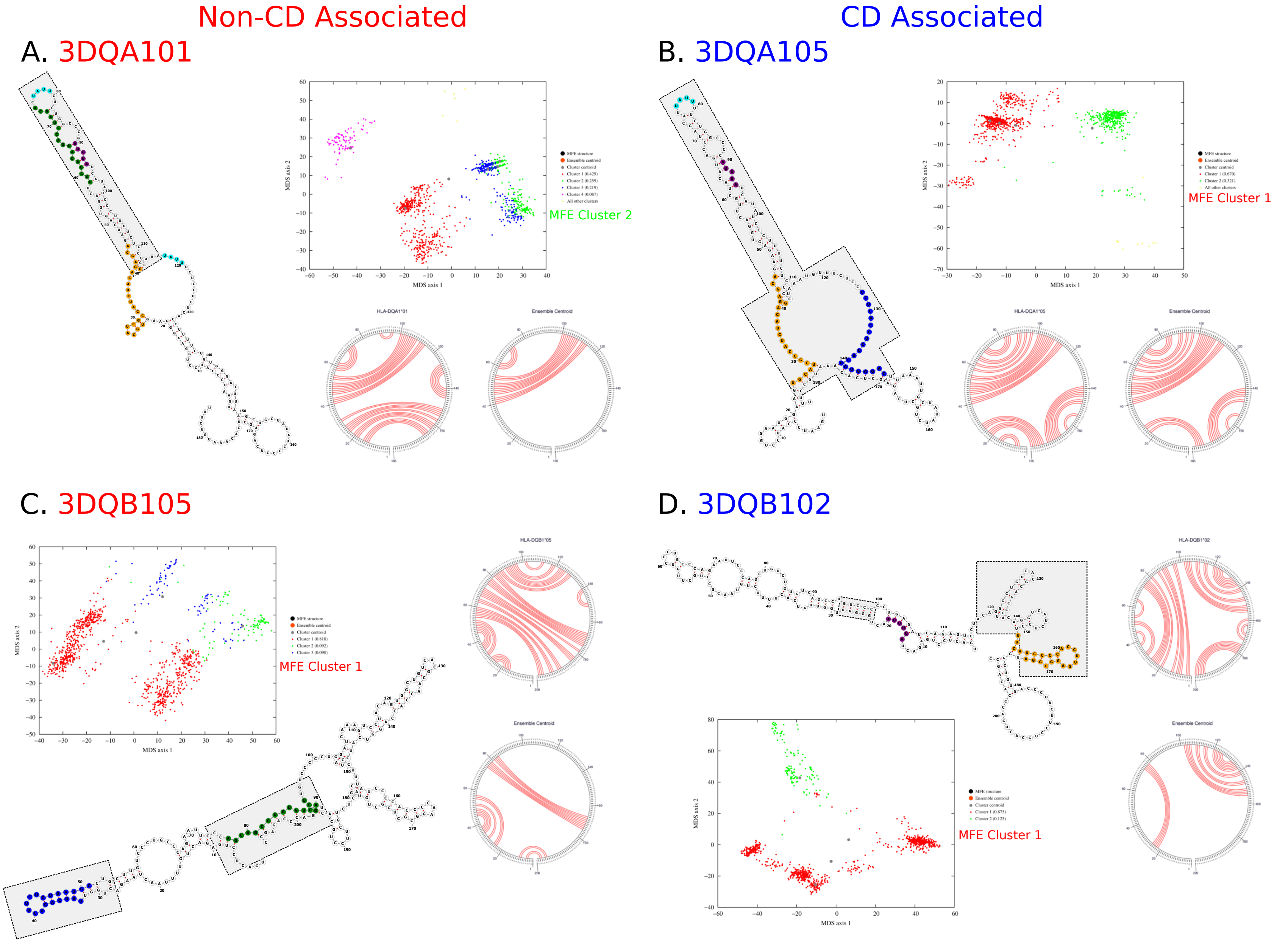

### Supplemental Figure 1

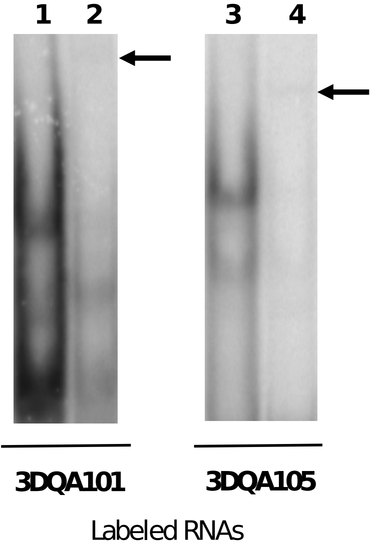
