## Supplemental Figure 3 for "Tristetraprolin/ZFP36 regulates the turnover of autoimmune-associated HLA-DQ mRNAs"

**A.**

```
3DQA101     UGAAUCCCAUCCUGGAAGGGAA-GUGCAUCGCCAUCUACAGGAGCAGAAGAGUGGACUUG
3DQA105     UGAAUCCCAUCCUGGAAUGGAA-GUGCAUCGCCAUCUACAGGAGCAGAAGAGUGGACUUG
H2-Aa-204   ---GUCACACCCUGGAAAGGAAGGCGUGUGUCCCUCUUCAGGGAAGAAGUGGUGUGCUGG

3DQA101     CU-ACAUGACCUAGCACUAUUCUCUGGCCCGAUUUAUCAUAUCCCUUUUCUCCUCCAAAU
3DQA105     CU-ACAUGACCUAGCAUUAUUUUCUGGCCCCAUUUAUCAUAUCCCUUUUCUCCUCCAAAU
H2-Aa-204   GUGACCUGGCACAGUG-UGUUUUCUGGACCAAUUCAUGGUGUUCUUUCUCUUCUUCAAGU

3DQA101     AUUUCUCCUCUCACCUUUUCUGUGGGACUUAAGCUGCUAUAUCCCCUCAGAGCUCACAAA
3DQA105     GUUUCUCCUCUCACCUCUUCUGUGGGACUUAAAUUGCUAUAUCUGCUCAGAGCUCACAAA
H2-Aa-204   GACCCCCAACUUGCUUUUCUCUUGACCCUGAGGCUGU----CCCUCUCACAGCUCACACA

3DQA101     UGUCUUU
3DQA105     UGCCUUU
H2-Aa-204   C-CCUUG
```

**B.**

| | 3DQA101 | 3DQA105 | H2-Aa-204 | GC |
|---|---|---|---|---|
| 3DQA101 | | 94.1 | 57.8 | 0.46 |
| 3DQA105 | 174/185 | | 57.8 | 0.44 |
| H2-Aa-204 | 108/187 | 108/187 | | 0.52 |

**C.**

```
3DQB102     UGACUCCUGAGACUAUUUUA-ACUGGGA---UUGGUUAUCACUUUUCUG---UAACGCCU
3DQB105     UGACUCCUGAGACUGUUUUA-ACUAAGA---CUGGUUAUCACUCUUCUG---UGAUGCCU
H2-Ab1-201  UGACUCAGUUGACUGUCUCAGACUGUAAGACCUGAAUGUCUCUGCUCCGAAUUCCUGACU

3DQB102     GCUUGUC--CCUGCC---------CAGAAUUC--CCAGCUGUCUGUGU-----CAGCCUG
3DQB105     GCUUGUC--CCUGCC---------CAGAAUUC--CCAGCUGCCUGUGU-----CAGCUUG
H2-Ab1-201  GCCAGUCCACCUGCCACUCCGACUCAGAGUCUAGCAUGGUACUAUUGUAUCCACCACCUC

3DQB102     UCCCCCUGAGAUC-AGAGUCCUACAG----UGGCUGUCACGCAGCCACCAGGU-------
3DQB105     UCCCCCUGAGAUC-AAAGUCCUACAG----UGGCUGUCACGCAACCACCAGGU-------
H2-Ab1-201  AGCUCUUGUGAUCUGGAGUCCCCAGUCUCUGUCUGUAGCUCUGCUCCUUGGUGAUUCCA

3DQB102     -------CAUCUCCUUUCAUCCCCACCUUGAGGCGGAUGGCUGUGACCC-----------
3DQB105     -------CAUCUCCUUUCAUCCCCACCCCAAGGCG-CUGGCUGUGACUC-----------
H2-Ab1-201  GAGACUCCAUCUGUGUACAGCCGCACCCCAGGCUUUCUGUCUUCGAUCUGUAGUAAACCA

3DQB102     -----UACUU-------CCUGCACU------------GACCCACAGCC
3DQB105     -----UGCUU-------CCUGCACU------------GACCCAGAGCC
H2-Ab1-201  AUGUAUGCUUAUCCCCACCUAGAUUACAAAUAAACGAGACUCAGACUC
```

**D.**

| | 3DQA102 | 3DQA105 | H2-Ab1-201 | GC |
|---|---|---|---|---|
| 3DQA102 | | 92.3 | 43.4 | 0.55 |
| 3DQA105 | 193/209 | | 45.8 | 0.55 |
| H2-Ab1-201 | 125/288 | 132/288 | | 0.50 |
